# Ploidy determines the consequences of antifungal-induced mutagenesis in Candida albicans, a human fungal pathogen

**DOI:** 10.1101/686881

**Authors:** Ognenka Avramovska, Meleah A. Hickman

## Abstract

Organismal ploidy state and environmental stress impact the mutational spectrum and the mutational rate. The human fungal pathogen *Candida albicans*, serves as a clinically relevant model for studying the interaction between eukaryotic ploidy and stress-induced mutagenesis. In this study, we compared the rates and types of genome perturbations in diploid and tetraploid *C. albicans* following exposure to two classes of antifungal drugs, azoles and echinocandins. We measured mutations at three different scales: point mutation, loss-of-heterozygosity (LOH), and genome size changes in cells treated with fluconazole and caspofungin. We find that caspofungin induced higher rates of mutation than fluconazole, likely an indirect result from the stress associated with cell wall perturbations rather than an inherent genotoxicity. Furthermore, we found disproportionately elevated rates of LOH and genome size changes in response to both antifungals in tetraploid *C. albicans* compared to diploid *C. albicans*, suggesting that the magnitude of stress-induced mutagenesis results from an interaction between ploidy state and the environment. These results have both clinical and evolutionary implications for how fungal pathogens generate mutations in response to antifungal drug stress, and may facilitate the emergence of antifungal resistance.

## Introduction

Mutations are the source material for adaptation. The mutational spectrum ranges from small-scale mutations, such as base substitutions and indels, to larger-scale mutational events such as gross chromosomal rearrangements and aneuploidy. Despite the presence of DNA repair mechanisms, all organisms incur mutations at a low level (Friedberg 2003); however, the rate of mutagenesis increases under stressful environments in both prokaryotes and eukaryotes (Tenaillon *et al.* 2004; Foster 2008; Petrosino *et al.* 2009; Forche *et al.* 2011; Maharjan and Ferenci 2017; Liu and Zhang 2019). For example, in yeast, the addition of inorganic salts, such as lithium chloride, increases the mutation rate by 3.5-fold (Liu and Zhang 2019). While stress increases mutational rates, it also shifts the mutational spectrum. Nutrient depletion, including phosphorus and nitrogen deficiencies, not only increase the mutation rate by seven-fold in *E. coli* but the types of mutations shift from indels to base substitutions (Maharjan and Ferenci 2017). Taken together, stressful environments can induce mutagenesis to provide increased genetic variation for natural selection to ultimately act upon.

While environmental stress impacts the rates and types of mutation, it is not the only factor that influences the mutational landscape. Ploidy, the number of sets of chromosomes an organism has, is an important determinant of the mutational spectrum available to an organism (Selmecki *et al.* 2015; Sharp *et al.* 2018). The human fungal pathogen, *C. albicans*, serves as a clinically-relevant model for studying the interaction between eukaryotic ploidy and stress-induced mutagenesis because of two key properties: the mutational rate and spectrum shifts extensively when *C. albicans* is exposed to stress (Forche *et al.* 2011) and *C. albicans* exists across a range of ploidy states – from haploid to polyploid (Rustchenko 2007; Selmecki *et al.* 2010; Hickman *et al.* 2015; Todd *et al.* 2017)

The ploidy state in which *Candida albicans* is isolated impacts its allelic phenotypes and genome stability. For example, *C. albicans* haploids have been isolated from a variety of *in vitro* stressors, such as the halo of a common antifungal susceptibility test and are mostly transient. They serve as a mechanism by which *C. albicans* can purge deleterious mutations, as the homozygous state incurs a fitness cost (Hickman *et al.* 2013). *C. albicans* is most-frequently isolated as a heterozygous diploid state (Jones *et al.* 2004; Braun *et al.* 2005; Abbey *et al.* 2011; Muzzey *et al.* 2013). In the diploid state, *C. albicans* frequently generates gross-chromosome rearrangements and maintains these large-scale mutations in laboratory and clinical isolates (Selmecki *et al.* 2009, 2010; Ford *et al.* 2015; Todd *et al.* 2017, 2019). Interestingly, diploid *C. albicans* undergo loss-of-heterozygosity (LOH) events 1000-fold more frequently than point mutations (Forche *et al.* 2011). Tetraploid *C. albicans* can be isolated from patients (Suzuki *et al.* 1986; Legrand *et al.* 2004; Abbey *et al.* 2014) and are generated in the laboratory through diploid-diploid mating or endoreplication (Forche *et al.* 2008). Tetraploids are pseudo-stable and undergo concerted chromosome loss to generate population heterozygosity through chromosome reassortment, aneuploidy, and SPO11-dependent recombination (Hull *et al.* 2000; Magee and Magee 2000; Bennett and Johnson 2003; Forche *et al.* 2008; Hickman *et al.* 2015). Gross-chromosomal rearrangements and loss-of-heterozygosity is 30-fold more frequent in tetraploid *C. albicans* compared to its diploid state (Hickman *et al.* 2015). Taken together, the mutational spectrums of haploid, diploid and tetraploid *C. albicans* are distinct, in accordance to other polyploid eukaryotes (Mayer and Aguilera 1990; Pavelka *et al.* 2010; Selmecki *et al.* 2015; Sharp *et al.* 2018).

*Candida albicans* is isolated in 40-60% of human invasive fungal infections (Pfaller and Diekema 2007, 2010; Pfaller *et al.* 2008; Pfaller 2012; Perfect 2017), and therefore, encounters diverse stressors within a human host. *In vitro* stress-induced mutagenesis studies show that stressors, such as febrile temperaturee (39°), elevate the rate of LOH and increase the rate of whole-chromosome rearrangements by ∼5-fold (Forche *et al.* 2011). *C. albicans* also encounters antifungal stressors, including azoles, which target the cell membrane, and echinocandins, which target the inner cell wall, because they are most commonly used to treat *C. albicans* infection (Pappas *et al.* 2016; Perfect 2017). Antifungal drugs may be mutagenic, as fluconazole exposure increases the rate of LOH in *C. albicans* (Forche *et al.* 2011) and induces aneuploidy (Harrison *et al.* 2014), aneuploidy is commonly with azole-resistant clinical isolates (Selmecki *et al.* 2006, 2009)With only a limited number of antifungals that can effectively treat these infections (Pappas *et al.* 2016), a deeper understanding of drug-induced mutagenesis is vital.

Unlike bacteria, which rapidly acquires drug resistance through plasmid transfer and horizontal gene transfer, resistance-associated mutations in *C. albicans* only arises through *de novo* events. Fluconazole resistance rates range from 1-86% in *Candida* species (Tsay *et al.* 2017; Pfaller *et al.* 2019) and can be due to copy number variation, homozygosis of drug-resistant alleles (Pfaller 2012; Ford *et al.* 2015; Morschhäuser 2016) and aneuploidy, including the formation of isochromosome 5L (Coste *et al.* 2006; Selmecki *et al.* 2006, 2009). In contrast, resistance to caspofungin, a commonly used echinocandin, is limited to single point mutations in *FKS1*, the gene encoding the drug target of echinocandins, a glucan synthase enzyme, (Pfaller 2012; Sanguinetti *et al.* 2015) While drug-induced mutagenesis has been studied in diploid *C. albicans* exposed to fluconazole, not much is known about how ploidy impacts this phenomena and how other clinically relevant antifungal drugs impact *C. albicans* genome stability. Though these dynamics have yet to be investigated, understanding them has implications for the evolution of drug-resistance in this common human opportunistic fungal pathogen.

In this study, we asked whether diploid and tetraploid *C. albicans* generate the same type of genomic perturbations when exposed to two classes of antifungal drugs. To address this question, we measured LOH rates and genome size changes in both diploid and tetraploid *C. albicans* exposed to fluconazole and to caspofungin. We found that regardless of ploidy, caspofungin was more mutagenic, through an indirect mechanism likely a result of cell wall stress, rather than genotoxicity. Additionally, we found that in the tetraploid state, *C. albicans* showed elevated rates of LOH and genome size changes in response to both antifungals compared to diploid *C. albicans*. These findings suggest that an interaction between ploidy state and the stress environment impacts the magnitude of stress-induced mutagenesis. These results have evolutionary and clinically relevant implications for how fungal pathogens generate adaptive mutations and how these mutations facilitate antifungal resistance.

## Results

### Antifungal drug susceptibility is similar for diploids and tetraploids

First we assessed if antifungal drug susceptibility differs between diploid and tetraploid *C. albicans*, and determined the minimum inhibitory concentration (MIC) of fluconazole (FLU) and caspofungin (CAS) for isogenic diploid and tetraploid *C. albicans*. We did not obsereve any differences in drug susceptibility between the two strains and measured the MIC for fluconazole to be ∼1 µg/mL and caspofungin to be <0.25 µg/mL (Fig. S1A & B). To determine the cell viability of these two ploidy states after antifungal drug exposure, we measured colony forming units (CFUs) following 24-hour exposure to 1 and 10 µg/mL fluconazole and 0.25 and 2.5 µg/mL caspofungin, representing 1× and 10× the MIC. While we do not detect significant reduction in the number of CFUs following exposure to fluconazole compared to the no drug treatment, we do see that exposure to fluconazole substantially slows diploid and tetraploid growth rates, particularly at high concentrations (S1C & S1D). We find that exposure to caspofungin reduces the number of viable CFUs more severely than fluconazole, regardless of ploidy (Fig. 1A & B). Interestingly, the low concentration of caspofungin inhibited cell growth to a greater degree than higher concentrations for both diploid and tetraploids *C. albicans*, a phenomenon coined the “the paradoxical effect of echinocandins” (Wagener and Loiko 2017). When we compare CFUs between tetraploid to diploid *C. albicans*, we observe a 50% reduction in tetraploids in the no drug treatment and detect a similar reduction in fluconazole, regardless of concentration (Fig. 1C). In contrast, both doses of caspofungin resulted in >90% reduction in tetraploid CFUs compared to diploid CFUs, suggesting that the tetraploid *C. albicans* is more sensitive to caspofungin than the diploid state.

**Figure 1:**
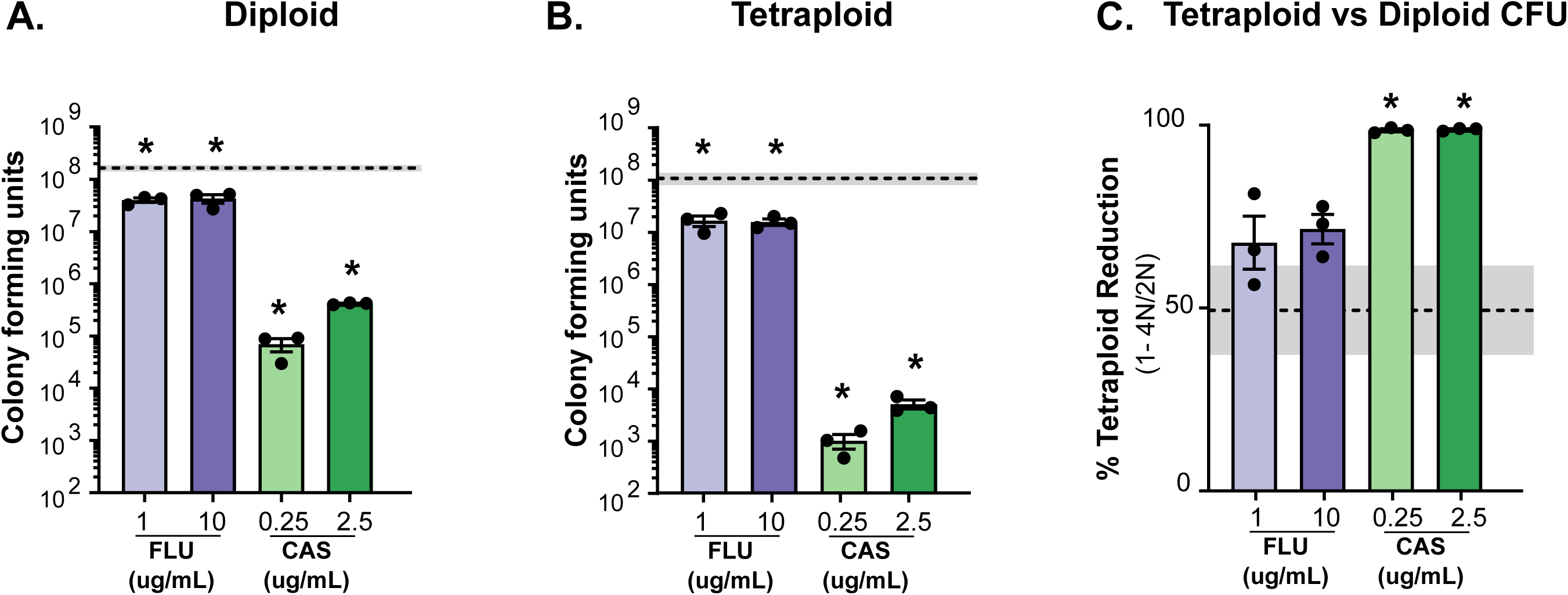
Diploids and Tetraploids are sensitive to antifungals. **A)** Diploid colony forming units following 24 h of exposire with no drug (dashed line). 1 ug/mL or 10 ug/mL fluconazole (‘FLU’, light and dark purple bars), 0.025 ug/mL or 2.5 ug/mL caspofungin (‘CAS’, light and dark green bars). The bars represent mean of at least 3 indepedent experiments (black symbols) and the error bars are +/-SEM. The dashed line and shaded box represent the mean, +/-SEM of CFUs obtained in no drug treatment. Asterisks indicate the drug treatments that differ significantly from no drug treatment (*, p <0.05; ** p < 0.01; unpaired Mann-Whitney U-tests). **B)** Tetraploid colony forming units. Analysis was performed same as A. **C)** The percent reduction of tetraploid CFUs relative to diploid CFUs was determined. Data is displayed and analyzed similarly to (A).

### Tetraploid mutation rates are differentially impacted by antifungal drugs compared to diploids

Given that short-term exposure impacts cell viability and growth of diploid and tetraploid *C. albicans*, we next wanted to investigate if antifungal drug exposure increased mutations rates. Since mutations can range from small scale, such as single nucleotide mutations, to large scale, such as gene conversion events and aneuploidy, we measured both the rates of point mutations using a *his4-G929T* (on Chr4) reversion assay (Forche *et al.* 2011) as well as a loss-of-heterozygosity (LOH) at the *GAL1* locus on Chr1 in diploid and tetraploid *C. albicans* following exposure to low and high concentrations of fluconazole and caspofungin. The reversion rate in the no drug treatment was extremely low for both diploid (∼4 × 10^10^ events/cell division) and tetraploid (∼12 × 10^10^ events/cell division) strains, albeit the mutations rate is slightly higher in tetraploid *C. albicans*. Exposure to antifungal drugs either did not change the mutation rate, or only modestly increased it (Table S1). For example, exposure to the high concentrations of fluconazole and caspofungin only increased the tetraploid mutation rate by two-fold. It should be noted that the rarity of revertants, coupled with the fungicidal nature of caspofungin, made it technically difficult to capture reversion events in all our experiments.

While point mutations are quite rare in *C. albicans*, large-scale genomic rearrangements occur more frequently and are easily detected (Forche *et al.* 2011; Hickman *et al.* 2015; Ene *et al.* 2018). To determine how exposure to antifungal drugs impacts the rate of large-scale rearrangements in diploid and tetraploid *C. albicans*, we measured the rate of *GAL1* loss-of-heterozygosity in both ploidy states. In diploid *C. albicans*, both concentrations of caspofungin significantly increased the rate of LOH compared to the no drug treatment, but exposure to fluconazole did not (Fig. 2A). In contrast, in tetraploid *C. albicans*, the rate of LOH was significantly higher in both fluconazole and caspofungin treatments compared to the no drug treatment (Fig. 2B). However, the degree to which the antifungal drugs elevated the rate of LOH differed; exposure to either concentration of fluconazole increased the LOH rate by ∼3-fold, whereas exposure to caspofungin increased the LOH rate by ∼100-fold compared to the no drug treatment.

**Figure 2:**
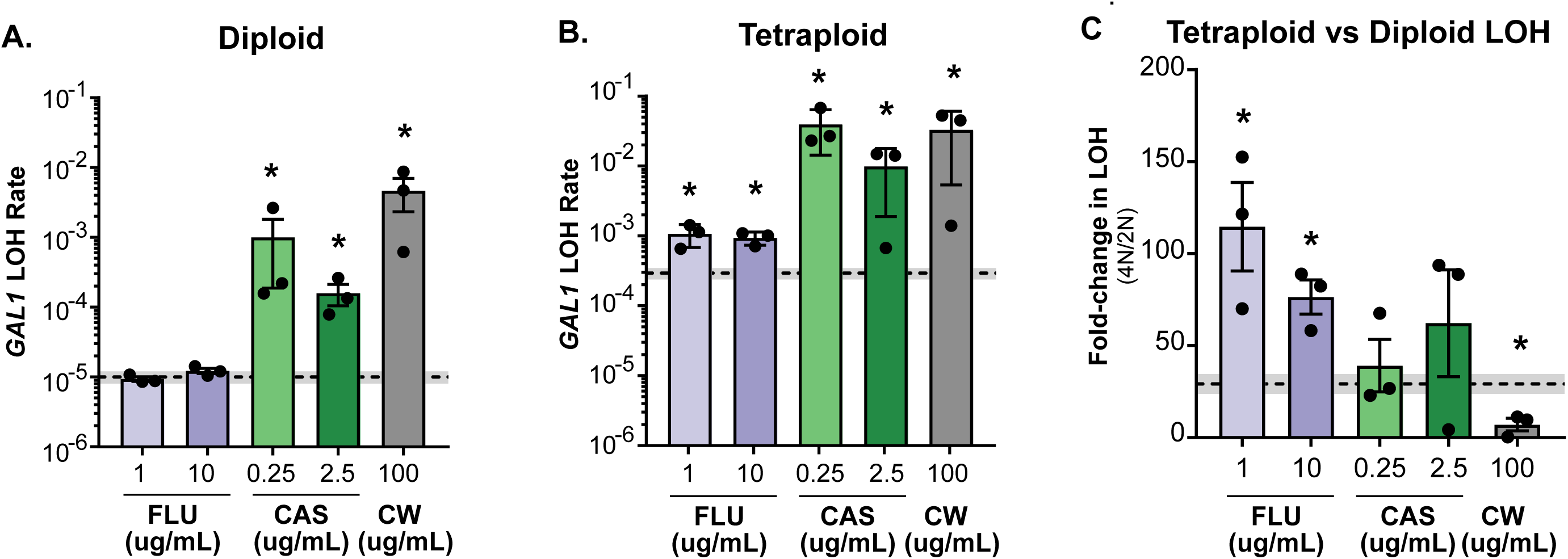
Ploidy- and antifungal drug-specific impacts on LOH in *C. albicans*. **A)** Diploid *GAL1* LOH rate after 24 h exposure to treatments with no drug (dashed line), 1 ug/mL or 10 ug/mL fluconazole (‘FLU’, light and dark purple bars, respectively), 0.25 ug/mL or 2.5 ug/mL caspofungin (‘CAS’, light and dark green bars, respectively), and the cell wall damaging agent, calcofluor white (‘CW’. grey bar). The bars represent the mean of at least three independent experiments (indicated with symbols) and the error bars represent the standard error of the mean (SEM). The dashed line and shaded box represents the mean *GAL1* LOH +/-one SEM of the no drug treatment. Asterisks indicate the drug treatments that differ significantly from the no drug treatment (* p < 0.05; unpaired Mann-Whitney U-tests). **B)** Tetraploid *GAL1* LOH. Analysis was performed the same as in (A). **C)** The fold-increase in *GAL1* LOH rate in tetraploid relative to diploid *C. albicans*. Data is displayed and analyzed similarly to (A).

We also observe significant differences in the overall rate of LOH between the diploid and tetraploid state (Fig. 2C). In the no drug treatment, the rate of LOH is 1.0 ×10^−5^ events/cell division in the diploid state (Fig. 2A, dashed line) compared to 2.9 ×10^−4^ events/cell division in the tetraploid state (Fig. 2B, dashed line) – an ∼30-fold difference, (Fig. 2C, dashed line), in accordance with previously published results (Hickman 2015). If there is not a ploidy-specific interaction with antifungal drugs on LOH rates, then we would expect to see a similar 30-fold increase of tetraploid LOH across the drug treatments. Indeed, this is pattern is observed in caspofungin when we compare the rate of LOH between the tetraploid and diploid, where the average fold-change in tetraploid LOH is ∼40 and is not statistically different than the no drug treatment (Fig. 2C, green bars). In contrast, both fluconazole concentrations have a >60-fold-change in tetraploid LOH, which is statistically different than the no drug treatment (Fig 2C, purple bars). Furthermore, we detect a significant interaction between *C. albicans* ploidy state and antifungal drugs on LOH rates (‘interaction’ p = 0.0019, two-way ANOVA) in addition to the individual impact from either ploidy (p = 0.0024, two-way ANOVA) or drug treatment (p = 0.0011, two-way ANOVA) (Fig. S2).

Regardless of ploidy state, we observed that exposure to caspofungin significantly increases LOH rates, a surprising result given that there is little supportive evidence for its genotoxicity, though some evidence suggesting it induces apoptosis in cells (Hao *et al.* 2013). To test whether the dramatic increase in LOH in caspofungin-treated *C. albicans* is directly related to is antifungal characteristics and potential genotoxicity or if the increased LOH is an indirect consequence of cell-wall stress, we exposed diploid and tetraploid *C. albicans* to 100ug/mL calcofluor white (Walker *et al.* 2008), a cell-wall perturbing agent at a concentration that inhibited growth to a similar degree as the low concentration of caspofungin (Fig. S3). We found that exposure to calcofluor white elevated the rate of LOH in both diploid (Fig. 2A) and tetraploid (Fig. 2B) *C. albicans* by 100-fold compared to the no drug treatment. This dramatic increase in LOH associated with calcofluor white is comparable to that of caspofungin and suggests that increased LOH is an indirect consequence of cell wall stress. Interestingly, when we compared the LOH rate between tetraploid and diploid *C. albicans*, there was only an 11-fold change between ploidy states, a statistically smaller difference than the no drug treatment (Fig. 2C, grey bar) and may indicate that the diploid state is more susceptible to cell wall stress than the tetraploid state.

### Genome size changes occur in response to antifungal drugs regardless of ploidy

Since exposure to antifungal drugs increased the rates of mutagenesis in either one or both ploidy states, we next examined whether genome size changes occur in response to these treatments. We measured total genome size of at least 80 single colony derivatives of diploid and tetraploid *C. albicans* after 24-hour exposure to antifungal drugs. We detect significant genome size changes across the set of single colonies isolated from both diploid (Fig. 3A) and tetraploid (Fig. 3B) *C. albicans* following exposure to nearly all antifungal drugs treatments compared to genome size distribution obtained from a similar number of isolates derived from the no drug treatment (Fig. 3A & B, grey shading). For diploid *C. albicans*, most genome size changes detected were modest increases in total DNA content. In particular, the high concentration of fluconazole and low concentration of caspofungin resulted in higher genome size in ∼8% and ∼20% of single colony isolates, respectively (Fig. 3C). Interestingly, these increases in genome size are likely transient, as we do not detect deviations from diploidy in isolates that have been exposed to antifungal drugs for longer time periods (Fig. S4A & C). In contrast to diploid *C. albicans*, tetraploid genome size changes we detect are predominantly losses in DNA content and are found in all drug treatments. The low concentration of fluconazole had the greatest impact, with nearly 40% of single colony isolates displaying small reductions in genome size (Fig. 3D). We also examined genome size changes associated with longer exposures to antifungal drugs (Fig. S4). While, we still detect a substantial number of single colony isolates derived from tetraploid *C. albicans* following caspofungin exposure that show genome size changes, instead of small the reductions in genome size that we detected at 24-hours, we now observe that most isolates have undergone massive genome reductions (Fig. S4B & D).

**Figure 3:**
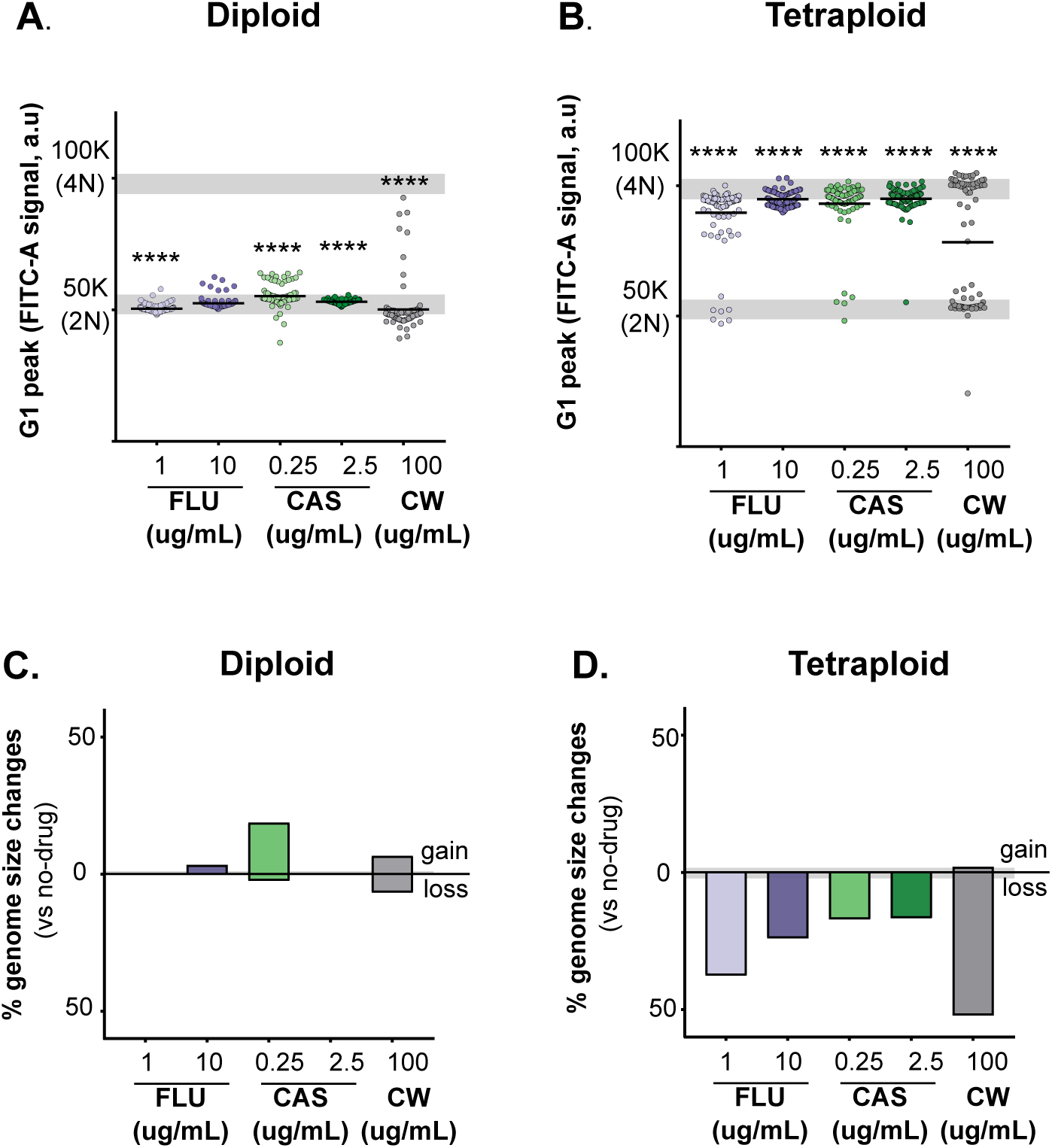
Antifungal drugs and cell wall stress induce genome size changes during short-term exposure. **A)** Total diploid genome size. Diploid genome size was measured after 24hrs of drug exposure using flow cytometry, with the G1 peak plotted in a.u (arbitrary units). The gray line represents the mean of the diploid no drug control, (n = 189) and the tetraploid no-drug control (n = 178) +/-1 SD. Each symbol represents the genome size average of 10,000 events from one culture (1 ug/mL FLU, n = 96, 10 ug/mL FLU, n = 89, 0.25 ug/mL CAS, n = 86, 2.5 ug/mL CAS, n = 93. 100 ug/mL CW, n = 90). The bold line represents the mean across all of the samples. Askrisks indicate statistical signifance compared to the no-drug control (gray bars), Mann-Whitney U-test. **B)** Same as A, using tetraploid strain (no stress, n = 87, 1 ug/mL FLU, n = 93, 10 ug/mL FLU, n = 96, 0.25 ug/mL CAS, n = 82, 2.5 ug/mL CAS, n = 96, 100 ug/mL CW, n = 96.) **C)** Quantification of diploid genome size changes. The percentage of diploids samples (from A) that show genome size changes after 24hrs was calculated as any sample that showed a genome size difference after drug treatment that was 2-standard deviations away from the mean of the no-drug diploid control. **D)** Quantification of tetraploid genome size changes. Calculated and presented same as in (C).

As caspofungin frequently induced large-scale genome size changes, we wanted to test whether these changes resulted from a direct genotoxicity of caspofungin or from cell wall stress, so we measured the frequency and magnitude of genome size changes when cells are exposed to calcofluor white. We found that 24-hour exposure to calcofluor white induced genome size changes in ∼14% of diploid derived single colony isolates; half of which are gains in DNA content and half of which are losses (Fig. 3C). However, genome size shifts are not detected at 120 hours of exposure, and may only exist transiently. In contrast, we observe that 50% of tetraploid derived single colony isolates reduced in genome size following 24 hour exposure to calcofluor white and ∼20% at 120 hours (Fig. 3D & Fig. S4D). Taken together, these results support a model that organismal ploidy state and the type of stress that the organism is exposed to, determines the effects and magnitude of the mutagenesis.

## Discussion

Stress-induced mutagenesis and organismal ploidy state both impact mutation rates and spectrum, but these two phenomena have often only been studied in isolation. Here, we investigated how antifungal drug stress impacts both mutation rates for small- and large-scale genomic events in diploid and tetraploid *C. albicans*. We find a significant interaction between ploidy and antifungal drug stress on genome instability (‘interaction’ p = 0.0160, two-way ANOVA, Fig S2). Specifically, we observe that caspofungin greatly increases genome instability in both diploid and tetraploid *C. albicans*, albeit to varying degrees, while fluconazole modestly elevates genome instability in tetraploids and not at all in diploids. Our findings support the model that organismal ploidy state impacts genome perturbations induced by specific stressors and this stress-induced increase in genetic variation has the potential for adaptive mutations to quickly arise.

While the genomic response to fluconazole has been well-studied in diploids and to a limited degree in tetraploids (Forche *et al.* 2011; Harrison *et al.* 2014; Hickman *et al.* 2015; Popp *et al.* 2017) the impact of caspofungin on *C. albicans* mutation rates is less understood, and is mostly investigated in regards to mutations conferring drug resistance. We found that the rate of LOH (Fig. 2) and rate of point mutations (Table S1) was substantially elevated in both diploid and tetraploid cells grown in the presence of caspofungin compared to no-drug and fluconazole. Furthermore, exposure to caspofungin elicits genome size changes in both diploid and tetraploids compared to no drug (Fig. 3, Fig. S4). These unexpected findings prompted us to test if caspofungin directly causes genotoxicity or if increased genome instability is an indirect result from stress to the cell wall. We found a similar increase in genome instability when cells were exposed to calcofluor white, which interferes with the proper construction of the fungal cell wall (Walker *et al.* 2008) (Figs. 2 and 3). This is consistent with the premise that the increased mutagenesis is an indirect consequence of cell wall stress and not due to a direct genotoxicity of caspofungin.

Exposure to different concentrations of caspofungin results in a ‘paradoxical effect’ on growth in several pathogenic fungi, including *C. albicans* and is a phenomenon in which cells can reconstitute growth in concentrations of caspofungin above the MIC, while remaining susceptible to caspofungin (Wagener and Loiko 2017). We observe a similar effect in our experiments for both diploids and tetraploids. We detect higher CFUs in 2.5 ug/mL caspofungin compared to 0.25 ug/mL caspofungin (Fig. 1) and lower LOH rates and frequency of ploidy change in the higher concentration. This suggests that low concentrations of caspofungin are more stressful than higher doses. There is evidence that *Candida albicans* restructure its cell wall in response to higher doses of caspofungin in order to better mediate this stress (Walker *et al.* 2008; Wagener and Loiko 2017).This caspofungin-mediated cell wall restructuring, in conjunction with our observed reduced genome instability at high caspofungin concentrations is consistent with the premise that high concentrations of caspofungin are less stressful and elicit weaker stress-induced mutagenesis than lower caspofungin concentrations.

While exposure to caspofungin elevated rates of mutagenesis in both diploids and tetraploids, the degree to which these ploidy states are impacted is different. Tetraploid genomes are inherently more unstable than diploid genomes (Hickman *et al.* 2015; Gerstein *et al.* 2017) and we also observe ∼30-fold increase in the rate of LOH in tetraploids compared to diploids in the absence of antifungal drugs (Fig 2). When comparing the rate of LOH between tetraploids and diploids across the stressors, we see a significantly higher fold-increase (compared to 30-fold in no-drugs) upon exposure to both caspofungin and fluconazole. This effect is particularly exaggerated in fluconazole, where the LOH increases by 3-fold in tetraploids but does not change diploids. We also see substantially more genome size changes associated with tetraploidy compared to diploidy regardless of how long cells are exposed to stressors. Most drastically, 70% of tetraploid isolates reduced to ∼diploid in size following five days of caspofungin and 30% due to calcofluor white exposure, but we did not detect any large scale changes in diploids under those conditions and timeframe (Fig S4C & D). While we detected rampant genome size changes in tetraploids during short exposures to fluconazole and calcofluor white, we did not see these changes persist at longer timeframes. While the overall frequency and scale of genome size changes in diploids exposed to fluconazole was lower, it was similar at 24-hour and five-day exposures. Together, these results suggest that there are interactions between ploidy and antifungal drugs that govern genome instability in *C. albicans*.

Several recent studies have implicated a role for ploidy in facilitating resistance to fluconazole. For example, drug-resistant clinical isolates are frequently aneuploid (Selmecki *et al.* 2006a, 2009) or even polyploidy (Abbey *et al.* 2014). Whether these aneuploid drug-resistant clinical isolates result from mis-segregation events in a diploid cell or from parasexual ploidy reduction (Forche *et al.* 2008; Hickman *et al.* 2015) of tetraploid cells in the host remains unclear. However *in vitro*, tetraploids rapidly, yet transiently, arise in response to high doses of fluconazole by decoupling cell growth and DNA replication, leading to trimeric structures and defects in budding (Harrison *et al.* 2014). Tetraploids can also arise *in vitro* via mating and potentially combine drug-resistant alleles that arose independently in diploid cells to generate mating products with even greater increases in their fluconazole MIC, (Popp *et al.* 2019). Regardless of whether the mechanism of tetraploid formation is mediated by mating or mitotic defects, the very high degree of tetraploid genome instability, particularly under antifungal drug stress, can generate aneuploidy and may facilitate resistance to fluconazole. In *S. cerevisiae*, tetraploids adapt much rapidly to nutrient-limited environments (Selmecki *et al.* 2015) and in *C. albicans*, stress-induced ploidy transitions have been proposed as a strategy for accelerating adaptation in this opportunistic fungal pathogen (Berman and Hadany 2012). Here we show that tetraploids generate a more diverse repertoire of mutations and at higher frequencies than diploids, particularly in response to azole and echinocandin drugs, and likely facilitate the rapid emergence of antifungal drug resistance.

## Materials and Methods

### Yeast strains and media

The stains used in this study are listed in Table S1. MH297 was constructed by replacing the wildtype *HIS4* open reading frame with the dominant drug-resistant *NAT* gene by lithium acetate transformation. Transformants were selected on YPD containing 50 ug/mL NAT and replica plated to media lacking histidine to identify candidates whose wildtype *HIS4* genes was replaced. MH296 is the mating product of two *his4Δ:: NAT/his4-G929T* diploid strains. All strains were stored as glycerol stocks −80°C and maintained on YPD medium (1% yeast extract, 2% bactopeptone, 2% glucose, 1.5% agar, 0.004% adenine, 0.008% uridine) at 30°C. For the assays in which antifungals were used, 1 mg/mL fluconazole stocks were made from powder (ACROS Organics CAS#86386-73-4) and diluted into DMSO. 1 mg/mL caspofungin stocks (Sigma-Aldrich CAS#179463-17-3) and 10mg/mL calcofluor white stocks (Sigma-Aldrich CAS#4404-43-7) were also made from powder and diluted into ddH_2_0. Liquid yeast cultures were grown in casitone (0.9% bacto-casitone, 0.5% yeast extract, 1% sodium citrate, 2% glucose), with or without antifungals. Plating on synthetic complete media (SDC) was used for enumerating total CFUs following drug exposure (0.17% yeast nitrogen base without amino acids or ammonium sulfate, 0.5% ammonium sulfate, 2 sodium hydroxide pellets, 0.004% uridine, 0.2% synthetic complete media, 2% agar, 2% glucose).

### Drug susceptibility

The minimum inhibitory concentration (MIC) for each strain was performed as previous described (Rosenburg et. al 2018) with the following modifications. Single colonies were inoculated in liquid YPD and incubated at 30°C with shaking for ∼16 h. Cultures subsequently normalized to 1 OD in casitone and serially diluted (two, 10-fold serial dilutions with 100 µl culture : 900 µl ddH_2_0 (10^−1^ and 10^−2^). 200uL of the diluted cells were plated onto casitone plates containing 1% agar and left to dry for 10 minutes before placing standardized E-test strips (Biomeureix) of caspofungin (gradient 0.002 ug/mL – 32 ug/mL) and fluconazole (gradient 0.016 ug/mL – 256 ug/mL). Plates were incubated at 30°C and photographed after 24 h growth.

### HIS4 reversion

The rate of *his4-G929T* reversion was performed as previously described (Forche *et al* 2011) with the following modifications. Ten single colonies of MH296 and MH297 were inoculated into 6 mL liquid YPD and incubated at 30°C with shaking for 24 h. 1 mL of the overnight culture was added to 4 mL of the following casitione media: no drug, 1 ug/mL fluconazole, 10 ug/mL fluconazole, 0.25 ug/mL caspofungin, 2.5 ug/mL caspofungin, and 100 ug/mL calcofluor white. All cultures were subsequently grown at 30°C with shaking for 5 days. To determine the number of viable colonies, 500 uL of each culture was serially diluted and plated on 100 × 15mm SDC plates. To determine the number of *HIS* revertants, the remaining 4.5 mL of each culture was harvested by centrifugation, washed with H_2_O, resuspended in 300uL ddH20 and plated onto 150mm × 15mm SDC – His plates. Plates were incubated at 30°C for 48 h. The rate of reversion was determined by fluctuation analysis using Luria-Delbruck method (Luria and lbrück 1943). All reversion experiments were performed in triplicate. A minimum of 48 single colonies per strain/ploidy were picked from SDC plates and stored as glycerol stocks for subsequent flow cytometry analysis.

### GAL1 loss of heterozygosity

The rate of *GAL1* loss of heterozygosity (LOH) was performed as previously described (Hickman *et al* 2015) with the following modifications. Twelve single colonies of MH84 and MH128 were inoculated into 2 mL casitone in the presence or absence of drugs. The cultures incubated at 30°C for 24h and harvested by centrifugation, washed once with ddH_2_0 and resuspended in 1mL ddH_2_0. 100 uL of the appropriate dilutions, (10^−5^ for no drug, 10^−4^ for fluconazole and 10^−2^ for caspofungin and calcofluor white treated cultures, of each culture was plated onto SDC to determine total cell viability, and onto 2-deoxygalactose (10^−1^ for no drug, 10^−1^ for fluconazole and 10^0^ for caspofungin and calcofluor white treated cultures (2-DOG; 0.17% yeast nitrogen base without amino acids, 0.5% ammonium sulfate, 0.004% uridine, 0.004% histidine, 0.006% leucine, 0.1% 2-deoxygalactose, 3% glycerol) to select for cells that spontaneously lost *GAL1* during the previous overnight growth in the presence or absence of antifungal drugs. SDC CFUs plated on SDC were counted following 48 h, and CFUs plated on 2-DOG were counted following 72 h growth. Rates of LOH were determined by the method of the median (Lea and Coulson 1949). All LOH experiments were performed in triplicate. At least 48 single colonies per strain/ploidy were picked from SDC plates and stored as glycerol stocks for subsequent flow cytometry analysis.

### Flow cytometry

Flow cytometry analysis was performed as previously published (Hickman et al. 2015). Briefly, 200uL of cells in midlog-phase were harvested, washed with distilled water, and resuspended in 20uL of 50 mM Tris (pH 8):50 mM EDTA (50:50 TE). Cells were fixed with 95% ethanol and incubated at 4° overnight. Cells were washed twice with 50:50 TE, resuspended in 50µL of 1 mg/ml RNAse A and incubated at 37° for 1 h. Cells were then collected, resuspended in 50µL of 5 mg/ml Proteinase K, and incubated at 37° for 30 m. Cells were subsequently washed once with 50:50 TE, resuspended in 50µL SybrGreen (1:100 of dilution in 50:50 TE), (Lonza, CAT#12001-798, 10,000x concentrated) incubated at 4° overnight collected via centrifugation and resuspended in 150 µL 50:50 TE, briefly sonicated, and run on a LSRII machine with laboratory diploid (MH1) and tetraploid (MH2) strains serving as calibration and internal controls. To estimate the average G1 peak FITC intensity, the multi-Gaussian cell cycle model used (FloJoV10).

### Statistical analysis

Statistical analysis was performed using GraphPad Prism 8 software. Comparisons to no-stress were performed using non-parametric, unpaired, Mann-Whitney U-test.

---

**Figure S1:**
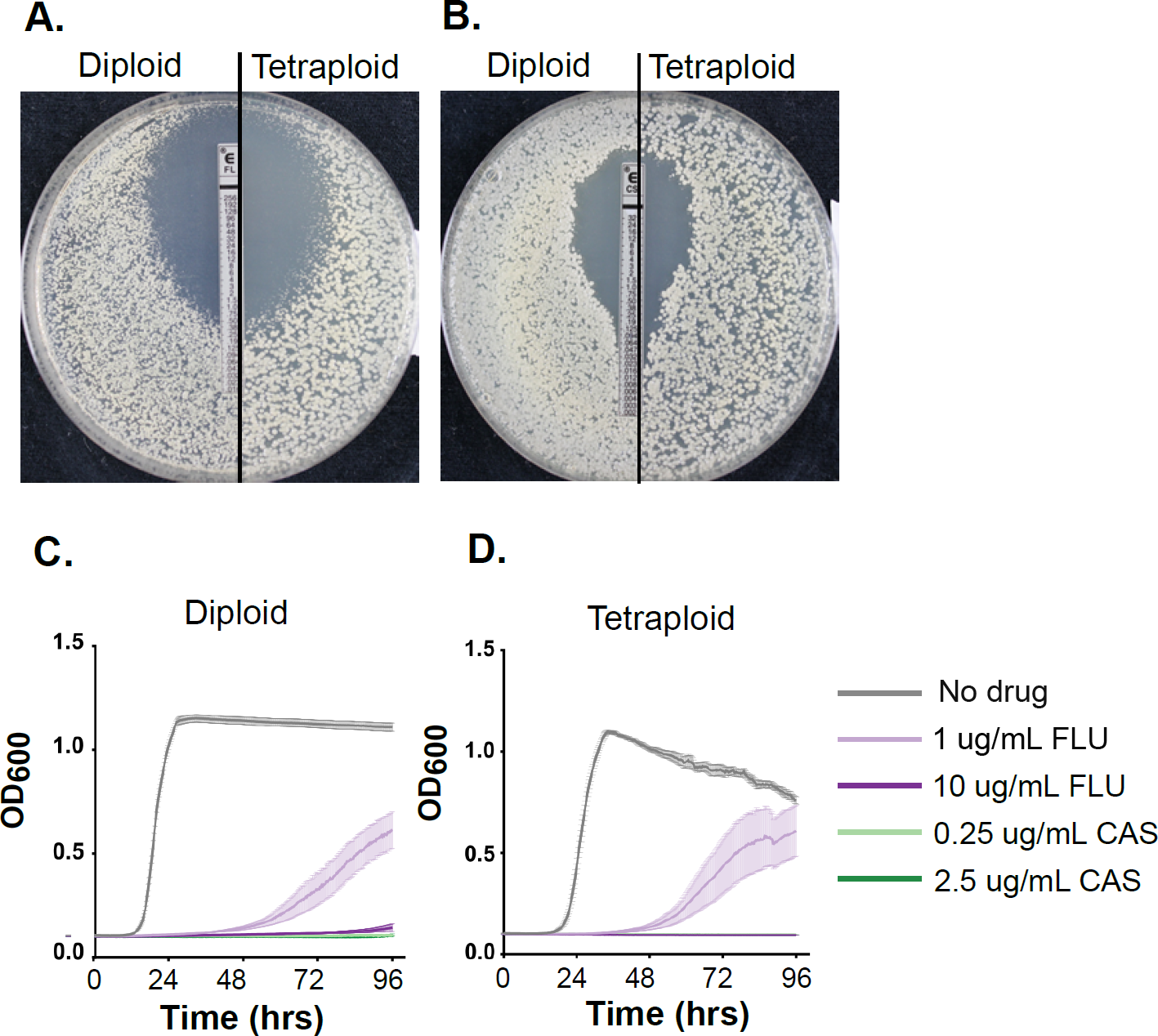
E-test and growth curves can quantify antifungal drug minimum inhibitory concentrations. **A)**. Diploid and tetraploid fluconazole MIC. The minimum inhibitory concentration (MIC) was determined using a standardized Biomerieux FLU e-test strip for 24hrs at 30°C. The size of the diameter in addition to the concentration of the strip where growth stops is indicative of the minimum inhibitory concentration. **B)** Diploid and tetraploid caspofungin MIC. The minimum inhibitory concentration (MIC) was determined using a standardized Biomerieux CAS e-test strip for 24hrs at 30°C. **C)** Diploid 96-hr frowth curve. The growth curves, shown as average diploid optical density (OD_600_) across at least 8 biological replicates is plotted over the course of 96hours. Each line represents a different treatment, gray – no drug (Casitone), lavender-1ug/mL FLU, dark purple-10ug/mL FLU, light green-0.25 ug/mL CAS and dark green-2.5 ug/mL CAS is plotted, with the average and error bars representing the SEM across the 8 replicates. **D)** Tetraploid 96-hr growth curve. Same as A, though testing at least 8 tetraploid isolates per condition.

**Supplemental Figure S2:**
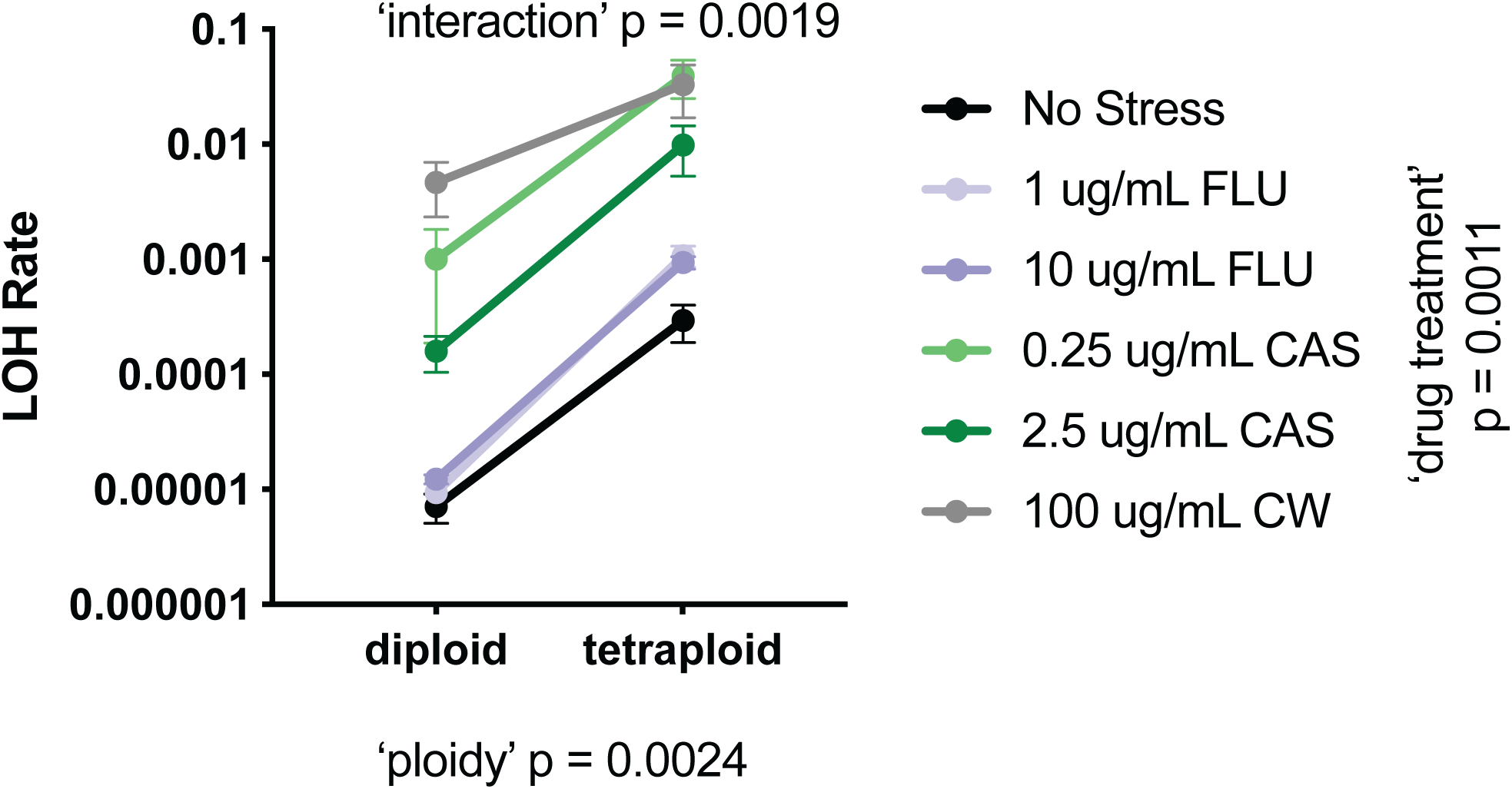
Two-Way ANOVA testing interaction between ‘drug treatment’ and ‘ploidy’ of Loss-of-heterozygosity measurements.

**Supplemental Figure 3:**
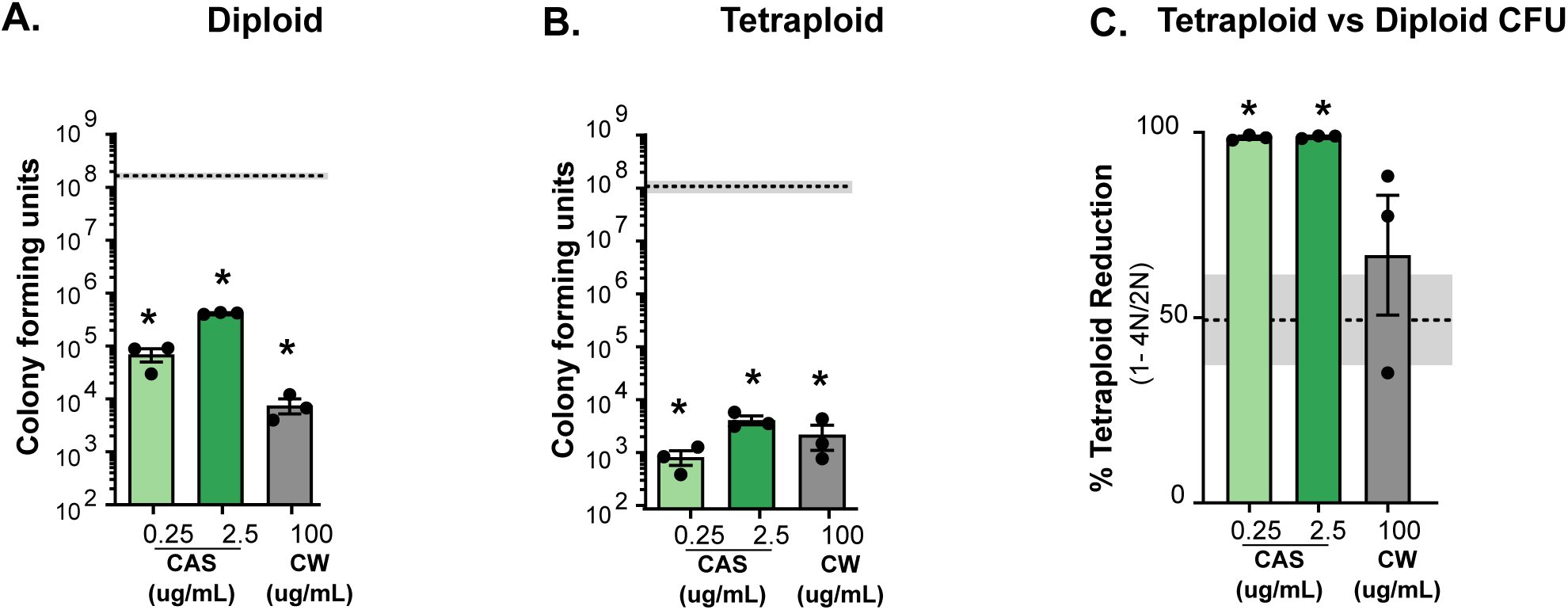
Diploids and Tetraploids are sensitive to antifungals. **A)** Diploid colony forming units following 24 h of exposure with no drug (dashed line). 0.025 ug/mL or 2.5 ug/mL caspofungin (‘CAS’, light and dark green bars) and 100ug/mL calcofluor while (‘CW’, gray). The bars represent mean of at least 3 indepedent experiments (black symbols) and the error bars are +/- SEM. The dashed line and shaded box represent the mean, +/- SEM of CFUs obtained in no drug treatment. Asterisks indicate the drug treatments that differ significantly from no drug treatment (*, p <0.05; ** p < 0.01; unpaired Mann-Whitney U-tests). **B)** Tetraploid colony forming units. Analysis was performed same as A. **C)** The percent reduction of tetraploid CFUs relative to diploid CFUs was determined. Data is displayed and analyzed similarly to (A).

**Supplemental Figure 4:**
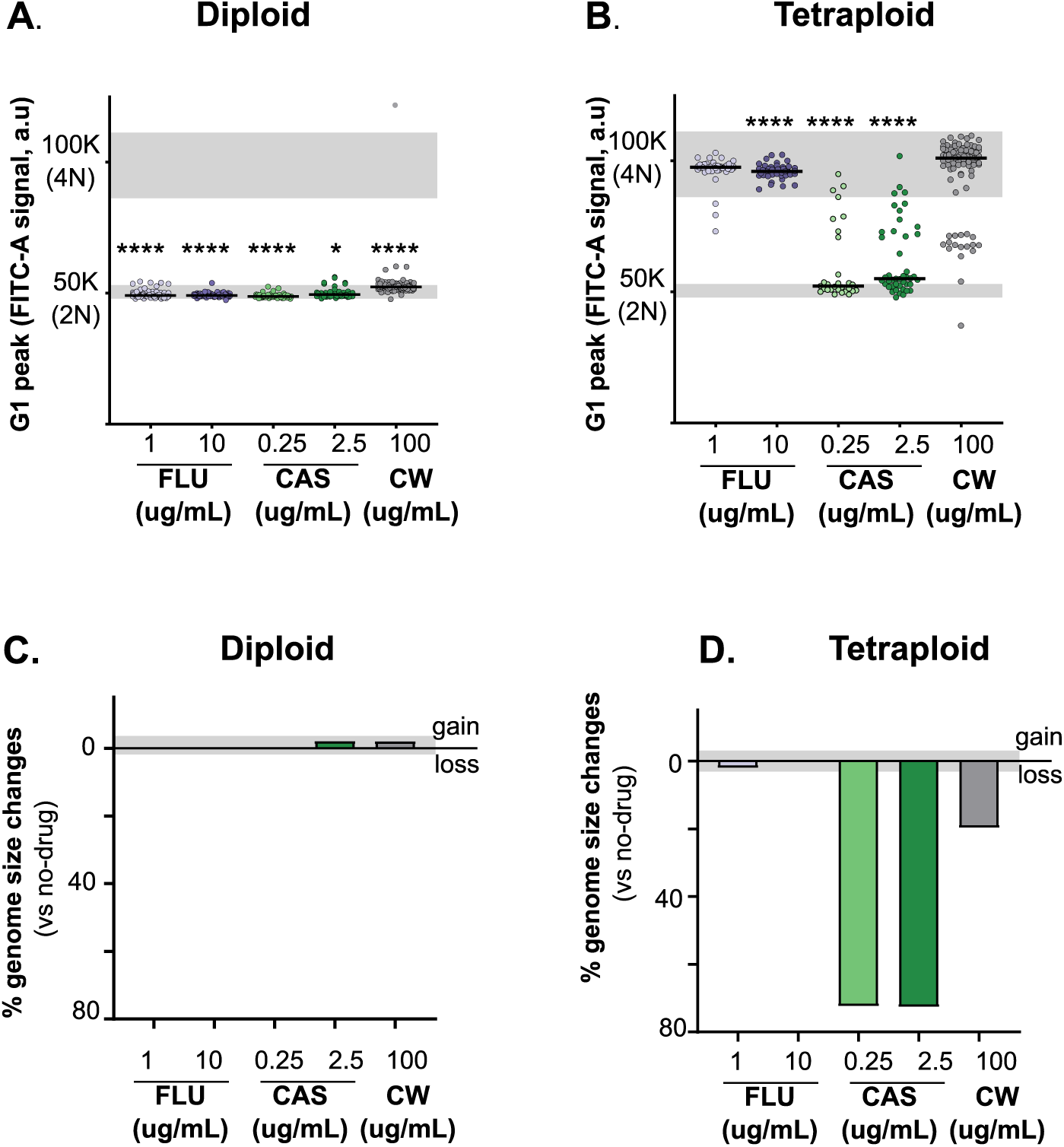
Caspofungin and calcoflour white induce large scale genome size changes in tetraploid *C. albicans* after 120 hrs of exposure. **A)** Diploid genome size was measured after 24hrs of drug exposure using flow cytometry, with the G1 peak plotted in a.u (arbitrary units). The gray lines represent the mean of the diploid no drug control, (n = 166), +/- 1SD and the tetraploid lab no-drug control, +/- 1SD. (n = 128). Each symbol represent the genome size average of 10,000 events from one culture (1 ug/ml FLU, n = 69, 10 ug/ml FLU, n = 72, 0.25 ug/ml CAS, n = 48, 2.5 ug/ ml CAS, n = 47, 100 ug/ml CW, n = 96). The bold line represents the mean across all of the samples. Asterisks indicate statistical significance compared to the no-drug diploid control, using Mann-Whitney U-test. **B)** Tetraploid genome size. Analysis and visualization same as (A), (1 ug/ml FLU, n = 47, 10 ug/ml FLU, n = 47, 0.25 ug/ml CAS, n = 39, 2.5 ug/ml CAS, n = 44, 100ug/ml CW, n = 66). **C)** Quantification of diploid genome size changes. The percentage of diploid samples from (A) that show and increase of decrease in genome size was calculated. An increase in genome size is any sample that had a G1 peak value (a.u) that was 2 SD above the diploid no-drug control. **D)** Quantification of tetraploid genome size changes. Analysis and visualization is same as (C).

**Table S1:** his4-G929T reversion rates in diploid and tetraploid *C. albicans.* The reversion rate was determined by fluctuation analysis using Luria-Delbruck Method (Luria S, Delbrück M 1943 and Spell and Jinks-Robertson, Molecular Biology 2004). The numbers represent the mean value across the experiments, while the number in parenthesis is the standard deviation across the independent trials. The ‘n’ value represents the number of independent experiments in which reversion events were observed.

| Strain | Ploidy | Reversion Rate (x 10<sup>-10</sup>) |  |  |  |  |
| --- | --- | --- | --- | --- | --- | --- |
|  |  | No drug | Fluconazole |  | Caspofungin |  |
|  |  |  | 1 ug/mL | 10 ug/mL | 0.25 ug/mL | 2.5 ug/mL |
| MH297 | diploid | 3.94 (±1.58) | 3.88 (±0.72) | 1.56 | n.d. | 63 |
|  |  | n=3 | n=3 | n=3 |  | n=1 |
| MH296 | tetraploid | 11.9 (±6.86) | 9.45 (±3.91) | 20.06 (±6.50) | 10.15 | 23.3 |
|  |  | n=3 | n=3 | n=3 | n=1 | n=1 |

**Table S2:**
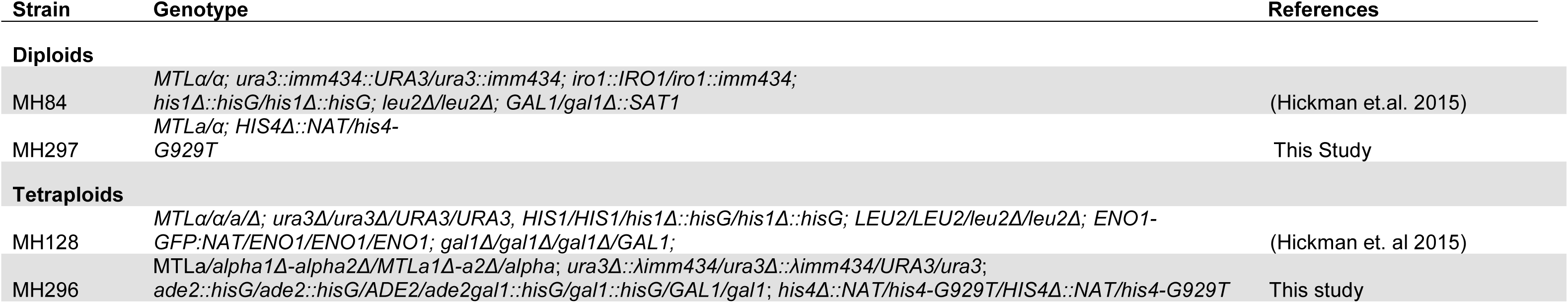
Strains used in this study.

## Literature Cited

Abbey D., M. Hickman, D. Gresham, and J. Berman, 2011 High-Resolution SNP/CGH Microarrays Reveal the Accumulation of Loss of Heterozygosity in Commonly Used Candida albicans Strains. G3 Genes Genomes Genetics 1: 523–530. https://doi.org/10.1534/g3.111.000885

Abbey D. A., J. Funt, M. N. Lurie-Weinberger, D. A. Thompson, A. Regev, et al., 2014 YMAP: a pipeline for visualization of copy number variation and loss of heterozygosity in eukaryotic pathogens. Genome Med 6: 100. https://doi.org/10.1186/s13073-014-0100-8

Bennett R. J., and A. D. Johnson, 2003 Completion of a parasexual cycle in Candida albicans by induced chromosome loss in tetraploid strains. Embo J 22: 2505–2515. https://doi.org/10.1093/emboj/cdg235

Berman J., and L. Hadany, 2012 Does stress induce (para)sex? Implications for Candida albicans evolution. Trends Genet 28: 197–203. https://doi.org/10.1016/j.tig.2012.01.004

Braun B. R., M. van het Hoog, C. d’Enfert, M. Martchenko, J. Dungan, et al., 2005 A Human-Curated Annotation of the Candida albicans Genome. Plos Genet 1: e1. https://doi.org/10.1371/journal.pgen.0010001

Coste A., V. Turner, F. Ischer, J. Morschhäuser, A. Forche, et al., 2006 A Mutation in Tac1p, a Transcription Factor Regulating CDR1 and CDR2, Is Coupled With Loss of Heterozygosity at Chromosome 5 to Mediate Antifungal Resistance in Candida albicans. Genetics 172: 2139–2156. https://doi.org/10.1534/genetics.105.054767

Ene I. V., R. A. Farrer, M. P. Hirakawa, K. Agwamba, C. A. Cuomo, et al., 2018 Global analysis of mutations driving microevolution of a heterozygous diploid fungal pathogen. Proc National Acad Sci 115: 201806002. https://doi.org/10.1073/pnas.1806002115

Forche A., K. Alby, D. Schaefer, A. D. Johnson, J. Berman, et al., 2008 The Parasexual Cycle in Candida albicans Provides an Alternative Pathway to Meiosis for the Formation of Recombinant Strains. Plos Biol 6: e110. https://doi.org/10.1371/journal.pbio.0060110

Forche A., D. Abbey, T. Pisithkul, M. Weinzierl, T. Ringstrom, et al., 2011 Stress Alters Rates and Types of Loss of Heterozygosity in Candida albicans. Mbio 2: e00129–11. https://doi.org/10.1128/mbio.00129-11

Ford C. B., J. M. Funt, D. Abbey, L. Issi, C. Guiducci, et al., 2015 The evolution of drug resistance in clinical isolates of Candida albicans. Elife 4: e00662. https://doi.org/10.7554/elife.00662

Foster P. L., 2008 Stress-Induced Mutagenesis in Bacteria. Crit Rev Biochem Mol 42: 373–397. https://doi.org/10.1080/10409230701648494

Friedberg E. C., 2003 DNA damage and repair. Nature 421: nature01408. https://doi.org/10.1038/nature01408

Gerstein A. C., H. Lim, J. Berman, and M. A. Hickman, 2017 Ploidy tug-of-war: Evolutionary and genetic environments influence the rate of ploidy drive in a human fungal pathogen. Evolution 71: 1025–1038. https://doi.org/10.1111/evo.13205

Hao B., S. Cheng, C. J. Clancy, and H. M. Nguyen, 2013 Caspofungin Kills Candida albicans by Causing both Cellular Apoptosis and Necrosis. Antimicrob Agents Ch 57: 326–332. https://doi.org/10.1128/aac.01366-12

Harrison B. D., J. Hashemi, M. Bibi, R. Pulver, D. Bavli, et al., 2014 A Tetraploid Intermediate Precedes Aneuploid Formation in Yeasts Exposed to Fluconazole. Plos Biol 12: e1001815. https://doi.org/10.1371/journal.pbio.1001815

Hickman M. A., G. Zeng, A. Forche, M. P. Hirakawa, D. Abbey, et al., 2013 The ‘obligate diploid’ Candida albicans forms mating-competent haploids. Nature 494: 55. https://doi.org/10.1038/nature11865

Hickman M. A., C. Paulson, A. Dudley, and J. Berman, 2015 Parasexual Ploidy Reduction Drives Population Heterogeneity Through Random and Transient Aneuploidy in Candida albicans. Genetics 200: 781–794. https://doi.org/10.1534/genetics.115.178020

Hull C. M., R. M. Raisner, and A. D. Johnson, 2000 Evidence for Mating of the “Asexual” Yeast Candida albicans in a Mammalian Host. Science 289: 307–310. https://doi.org/10.1126/science.289.5477.307

Jones T., N. A. Federspiel, H. Chibana, J. Dungan, S. Kalman, et al., 2004 The diploid genome sequence of Candida albicans. P Natl Acad Sci Usa 101: 7329–7334. https://doi.org/10.1073/pnas.0401648101

Lea D., and C. Coulson, 1949 The distribution of the numbers of mutants in bacterial populations. J Genet 49: 264. https://doi.org/10.1007/bf02986080

Legrand M., P. Lephart, A. Forche, F. C. Mueller, T. Walsh, et al., 2004 Homozygosity at the MTL locus in clinical strains of Candida albicans: karyotypic rearrangements and tetraploid formation†. Mol Microbiol 52: 1451–1462. https://doi.org/10.1111/j.1365-2958.2004.04068.x

Liu H., and J. Zhang, 2019 Yeast Spontaneous Mutation Rate and Spectrum Vary with Environment. Curr Biol. https://doi.org/10.1016/j.cub.2019.03.054

Luria S., and lbrück, 1943 Mutations of Bacteria from Virus Sensitivity to Virus Resistance. Genetics 28: 491–511.

Magee B., and P. Magee, 2000 Induction of Mating in Candida albicans by Construction of MTLa and MTLα Strains. Science 289: 310–313. https://doi.org/10.1126/science.289.5477.310

Maharjan R. P., and T. Ferenci, 2017 A shifting mutational landscape in 6 nutritional states: Stress-induced mutagenesis as a series of distinct stress input–mutation output relationships. Plos Biol 15: e2001477. https://doi.org/10.1371/journal.pbio.2001477

Mayer V. W., and A. Aguilera, 1990 High levels of chromosome instability in polyploids of Saccharomyces cerevisiae. Mutat Res Fundam Mol Mech Mutagen 231: 177–186. https://doi.org/10.1016/0027-5107(90)90024-x

Morschhäuser J., 2016 The development of fluconazole resistance in Candida albicans – an example of microevolution of a fungal pathogen. J Microbiol 54: 192–201. https://doi.org/10.1007/s12275-016-5628-4

Muzzey D., K. Schwartz, J. S. Weissman, and G. Sherlock, 2013 Assembly of a phased diploid Candida albicansgenome facilitates allele-specific measurements and provides a simple model for repeat and indel structure. Genome Biol 14: R97. https://doi.org/10.1186/gb-2013-14-9-r97

Pappas P. G., C. A. Kauffman, D. R. Andes, C. J. Clancy, K. A. Marr, et al., 2016 Clinical Practice Guideline for the Management of Candidiasis: 2016 Update by the Infectious Diseases Society of America. Clin Infect Dis 62: e1–e50. https://doi.org/10.1093/cid/civ933

Pavelka N., G. Rancati, J. Zhu, W. D. Bradford, A. Saraf, et al., 2010 Aneuploidy confers quantitative proteome changes and phenotypic variation in budding yeast. Nature 468: 321. https://doi.org/10.1038/nature09529

Perfect J. R., 2017 The antifungal pipeline: a reality check. Nat Rev Drug Discov 16: 603–616. https://doi.org/10.1038/nrd.2017.46

Petrosino J. F., R. S. Galhardo, L. D. Morales, and S. M. Rosenberg, 2009 Stress-Induced β-Lactam Antibiotic Resistance Mutation and Sequences of Stationary-Phase Mutations in the Escherichia coli Chromosome▿. J Bacterio l 191: 5881–5889. https://doi.org/10.1128/jb.00732-09

Pfaller M., and D. Diekema, 2007 Epidemiology of Invasive Candidiasis: a Persistent Public Health Problem. Clin Microbiol Rev 20: 133–163. https://doi.org/10.1128/cmr.00029-06

Pfaller M., D. Diekema, L. Ostrosky-Zeichner, J. Rex, B. Alexander, et al., 2008 Correlation of MIC with Outcome for Candida Species Tested against Caspofungin, Anidulafungin, and Micafungin: Analysis and Proposal for Interpretive MIC Breakpoints▿. J Clin Microbiol 46: 2620–2629. https://doi.org/10.1128/jcm.00566-08

Pfaller M. A., and D. J. Diekema, 2010 Epidemiology of Invasive Mycoses in North America. Crit Rev Microbiol 36: 1–53. https://doi.org/10.3109/10408410903241444

Pfaller M. A., 2012 Antifungal Drug Resistance: Mechanisms, Epidemiology, and Consequences for Treatment. Am J Medicine 125: S3–S13. https://doi.org/10.1016/j.amjmed.2011.11.001

Pfaller M. A., D. J. Diekema, J. D. Turnidge, M. Castanheira, and R. N. Jones, 2019 Twenty Years of the SENTRY Antifungal Surveillance Program: Results for Candida Species From 1997–2016. Open Forum Infect Dis 6: S79–S94. https://doi.org/10.1093/ofid/ofy358

Popp C., I. A. Hampe, T. Hertlein, K. Ohlsen, D. P. Rogers, et al., 2017 Competitive Fitness of Fluconazole-Resistant Clinical Candida albicans Strains. Antimicrob Agents Ch 61: e00584–17. https://doi.org/10.1128/aac.00584-17

Popp C., B. Ramírez-Zavala, S. Schwanfelder, I. Krüger, and J. Morschhäuser, 2019 Evolution of Fluconazole-Resistant Candida albicans Strains by Drug-Induced Mating Competence and Parasexual Recombination. Mbio 10. https://doi.org/10.1128/mbio.02740-18

Rustchenko E., 2007 Chromosome instability in Candida albicans. Fems Yeast Res 7: 2–11. https://doi.org/10.1111/j.1567-1364.2006.00150.x

Sanguinetti M., B. Posteraro, and C. Lass-Flörl, 2015 Antifungal drug resistance among Candida species: mechanisms and clinical impact. Mycoses 58: 2–13. https://doi.org/10.1111/myc.12330

Selmecki A., A. Forche, and J. Berman, 2006a Aneuploidy and Isochromosome Formation in Drug-Resistant Candida albicans. Science 313: 367–370. https://doi.org/10.1126/science.1128242

Selmecki A., A. Forche, and J. Berman, 2006b Aneuploidy and Isochromosome Formation in Drug-Resistant Candida albicans. Science 313: 367–370. https://doi.org/10.1126/science.1128242

Selmecki A. M., K. Dulmage, L. E. Cowen, J. B. Anderson, and J. Berman, 2009 Acquisition of Aneuploidy Provides Increased Fitness during the Evolution of Antifungal Drug Resistance. Plos Genet 5: e1000705. https://doi.org/10.1371/journal.pgen.1000705

Selmecki A., A. Forche, and J. Berman, 2010 Genomic Plasticity of the Human Fungal Pathogen Candida albicans▿. Eukaryot Cell 9: 991–1008. https://doi.org/10.1128/ec.00060-10

Selmecki A. M., Y. E. Maruvka, P. A. Richmond, M. Guillet, N. Shoresh, et al., 2015 Polyploidy can drive rapid adaptation in yeast. Nature 519: 349. https://doi.org/10.1038/nature14187

Sharp N. P., L. Sandell, C. G. James, and S. P. Otto, 2018 The genome-wide rate and spectrum of spontaneous mutations differ between haploid and diploid yeast. Proc National Acad Sci 115: 201801040. https://doi.org/10.1073/pnas.1801040115

Suzuki T., A. Rogers, and P. Magee, 1986 Inter- and intra-species crosses between Candida albicans and Candida guilliermondii. Yeast 2: 53–58. https://doi.org/10.1002/yea.320020104

Tenaillon O., E. Denamur, and I. Matic, 2004 Evolutionary significance of stress-induced mutagenesis in bacteria. Trends Microbiol 12: 264–270. https://doi.org/10.1016/j.tim.2004.04.002

Todd R. T., A. Forche, and A. Selmecki, 2017 Ploidy Variation in Fungi: Polyploidy, Aneuploidy, and Genome Evolution. Microbiol Spectr 5. https://doi.org/10.1128/microbiolspec.funk-0051-2016

Todd R. T., T. D. Wikoff, A. Forche, and A. Selmecki, 2019 Genome plasticity in Candida albicans is driven by long repeat sequences. Elife 8: e45954. https://doi.org/10.7554/elife.45954

Tsay S., R. M. Welsh, E. H. Adams, N. A. Chow, L. Gade, et al., 2017 Notes from the Field: Ongoing Transmission of Candida auris in Health Care Facilities — United States, June 2016– May 2017. Mmwr Morbidity Mortal Wkly Rep 66: 514–515. https://doi.org/10.15585/mmwr.mm6619a7

Wagener J., and V. Loiko, 2017 Recent Insights into the Paradoxical Effect of Echinocandins. J Fungi 4: 5. https://doi.org/10.3390/jof4010005

Walker L. A., C. A. Munro, I. de Bruijn, M. D. Lenardon, A. McKinnon, et al., 2008 Stimulation of Chitin Synthesis Rescues Candida albicans from Echinocandins. Plos Pathog 4: e1000040. https://doi.org/10.1371/journal.ppat.1000040

